# DYRK1A regulates the recruitment of 53BP1 to the sites of DNA damage in part through interaction with RNF169

**DOI:** 10.1101/327510

**Authors:** Vijay R. Menon, Varsha Ananthapadmanabhan, Selene Swanson, Siddharth Saini, Fatmata Sesay, Vasily Yakovlev, Laurence Florens, James A. DeCaprio, Michael P. Washburn, Mikhail Dozmorov, Larisa Litovchick

## Abstract

Human *DYRK1A* gene encoding Dual-specificity tyrosine (Y)- Regulated Kinase 1A (DYRK1A) is a dosage-dependent gene whereby either trisomy or haploinsufficiency result in developmental abnormalities. However, the function and regulation of this important protein kinase are not fully understood. Here we report proteomic analysis of DYRK1A in human cells that revealed a novel role of DYRK1A in the DNA double-strand break (DSB) repair signaling. This novel function of DYRK1A is mediated in part by its interaction with ubiquitin-binding protein RNF169 that regulates the choice between homologous recombination (HR) and non-homologous end joining (NHEJ) DSB repair. Accumulation of RNF169 at the DSB sites promotes homologous recombination (HR) by limiting the recruitment of the scaffold protein 53BP1 that promotes NHEJ by protecting the DNA ends from resection. Inducible overexpression of active, but not the kinase inactive, DYRK1A in U-2 OS cells inhibited accumulation of 53BP1 at the DSB sites in RNF169-dependent manner. Mutation of DYRK1A phosphorylation sites in RNF169 or pharmacological inhibition of DYRK1A using harmine decreased the ability of RNF169 to displace 53BP1 from radiation-induced DSB sites. In order to further investigate the role of DYRK1A in regulation of DNA repair, we used CRISPR-Cas9 mediated knockout of DYRK1A in human and mouse cells. Interestingly, knockout of DYRK1A also caused a defect in 53BP1 DSB recruitment that was independent of RNF169, suggesting that dosage of DYRK1A can influence the DNA repair processes through several mechanisms. U-2 OS cells devoid of DYRK1A displayed an increased DNA repair and HR efficiency, and showed a decreased sensitivity to the PARP inhibitor olaparib when compared to control cells. Given evidence of its altered expression in human cancers, DYRK1A levels could represent a significant determinant of the DNA damaging therapy response.

## Introduction

Gene dosage imbalance that involves regulatory genes, such as *DYRK1A*, could have dramatic consequences for the individual cells, tissues, organs or entire organisms (Birchler et al., 2005; Veitia and Potier, 2015). In case of *DYRK1A*, both gain and loss of one allele result in developmental abnormalities. Trisomy of a critical region on human chromosome 21 where the *DYRK1A* gene is located results in Down syndrome (DS) (Korbel et al., 2009; Ohira et al., 1997). Loss or intragenic deletion affecting one copy of the *DYRK1A* gene has been also recently recognized as a syndrome characterized by microcephaly and severe mental retardation (Bronicki et al., 2015; Ji et al., 2015). The requirement of the proper *DYRK1A* gene dosage for neurological development is conserved in evolution, as evident from genetic studies of its *Drosophila* ortholog *minibrain* (*mnb*) (Guimera et al., 1996; Tejedor et al., 1995). Furthermore, mouse models of the *Dyrk1a* trisomy recapitulate some of the DS phenotypes (Ahn et al., 2006; Altafaj et al., 2001; Tejedor and Hammerle, 2011). Homozygous deletion of *Dyrk1a* causes early embryonic lethality whereas Dyrk1a^+/−^ animals have reduced brain size as well as specific neurological and behavioral defects (Fotaki et al., 2004; Raveau et al., 2018). In order to explain these phenotypes, it is important to understand the function and regulation of DYRK1A.

DYRK1A belongs to the CMGC group of protein kinases that also includes cyclin-dependent kinases (CDKs), mitogen activated protein kinases (MAPKs), glycogen synthase kinases (GSKs), and CDK-like kinases (CLKs) (Aranda et al., 2010; Kannan and Neuwald, 2004). Functionally, DYRK1A is a dual-specificity protein kinase that regulates several protein substrates some of which are involved in control of the cell cycle and transcription including cyclin D1, p27, RNA polymerase II and LIN52 subunit of the DREAM repressor complex (Chen et al., 2013; Di Vona et al., 2015; Litovchick et al., 2011; Najas et al., 2015; Park et al., 2009; Soppa et al., 2014). DYRK1A preferentially phosphorylates protein substrates that match the consensus R-X(XX)-S-P where X is any amino acid (Becker and Sippl, 2011; Himpel et al., 2000) although some substrates such as cyclin D1 contain alternative phosphorylation sites (Chen et al., 2013; Soppa et al., 2014). In addition to these potential substrates, DYRK1A interacts with several proteins that may regulate its function or subcellular localization including DCAF7 and 14-3-3 (Alvarez et al., 2007; Kim et al., 2004; Miyata and Nishida, 2011; Ritterhoff et al., 2010). A recent study of the proteomic landscape of the CMGC kinases in HEK293T cells identified 24 cellular proteins specifically interacting with DYRK1A, including DCAF7 (Varjosalo et al., 2013). Furthermore, DYRK1A has been shown to interact with several viral proteins including adenovirus E1A and human papilloma virus E6 proteins, and alter their ability to transform host cells (Cohen et al., 2013; Komorek et al., 2010; Kuppuswamy et al., 2013; Subramanian et al., 2013).

Previously, we described a critical role of DYRK1A in the G0/G1 entry in human T98G glioblastoma cells by promoting the assembly of the DREAM transcription repressor complex (Guiley et al., 2015; Litovchick et al., 2011; Litovchick et al., 2007). Ectopic expression of DYRK1A suppressed proliferation of several human cell lines such as T98G and U-2 OS, but not HEK293T cells (Litovchick et al., 2011), suggesting that DYRK1A function could be influenced by cell-specific context. Therefore, we sought to characterize DYRK1A interacting proteins in T98G cells, using sensitive MudPIT proteomic analysis approach (Litovchick et al., 2011). Our analysis identified proteins that reproducibly and selectively co-precipitated with DYRK1A, including both previously reported and novel interactions. Here, we describe a novel role of DYRK1A in repair of DNA double-strand breaks (DSB) revealed through its interaction with the ubiquitin-binding protein, RNF169. Upon DNA damage, RNF169 accumulates at the DSBs and promotes homologous recombination repair (HRR) by restraining accumulation of 53BP1, a scaffolding protein associated with non-homologous end joining (NHEJ)-promoting factor, at the DSB sites (Chen et al., 2012; Panier et al., 2012; Poulsen et al., 2012). We found that DYRK1A regulates the recruitment of RNF169 and 53BP1 to the sites of DNA damage, and therefore the levels of DYRK1A in the cells can affect the choice of DNA repair pathway and the cell response to DNA-damaging drugs.

## Results

### MudPIT analysis of DYRK1A-interacting proteins

DYRK1A plays an essential role in cell cycle control in human T98G cells (Litovchick et al., 2011); therefore, we chose these cells for characterization of DYRK1A-interacting proteins using MudPIT MS/MS proteomic analysis (Florens and Washburn, 2006). HA-tagged DYRK1A was expressed in T98G cells (Figure 1A), purified using anti-HA affinity matrix and analyzed by MudPIT as previously described (Litovchick et al., 2011; Litovchick et al., 2007). Four biological replicates were analyzed for DYRK1A-HA pull-down samples along with 3 GFP-HA (control) samples, resulting in identification of 120 proteins (including DYRK1A) that were detected at least twice in the DYRK1A pull-down samples but not in the GFP controls (Table S1). Previous proteomic analysis of DYRK1A in HEK293 cells identified 24 interacting proteins, 14 of which were also detected in our study (Figure S1A) (Varjosalo et al., 2013). Furthermore, our analysis detected 51 proteins in 3 out of 4 DYRK1A pull-down repeats and 7 proteins including DYRK1A, DCAF7, FAM117A, FAM117B, LZTS2, RNF169 and TROAP, were identified in all biological replicates. These interacting proteins were also the most enriched in the samples and readily confirmed using reciprocal pull-down assays (Figures 1B, C and S1). Of note, average enrichment of DCAF7 in the immunoprecipitated samples (as shown by Normalized Spectrum Abundance Factor, or dNSAF (Sardiu et al., 2008)), was comparable to that of DYRK1A itself, indicating a potentially stoichiometric interaction (Figure 1C). Further bioinformatic analysis of the 51 DYRK1A-binding proteins revealed a complex network of interactions of factors involved in different cellular processes with notable enrichment of the mRNA processing, transcription and DNA damage response functional categories (Figures 1D and E). Interestingly, recent proteomic analysis of RNF169 also identified DYRK1A as one of the most enriched interacting proteins (An et al., 2017). RNF169 is a RING-domain ubiquitin-binding protein that plays a role in the DNA repair signaling by regulating 53BP1, a scaffold protein that plays a key role in the choice of the double strand break (DSB) DNA repair by promoting non-homologous end joining (NHEJ) and inhibiting homologous recombination (HR) (Chen et al., 2012; Panier and Boulton, 2014; Poulsen et al., 2012). Since the role of DYRK1A in DNA repair processes is not known, we chose to further characterize the interaction between DYRK1A and RNF169.

**Figure 1.**
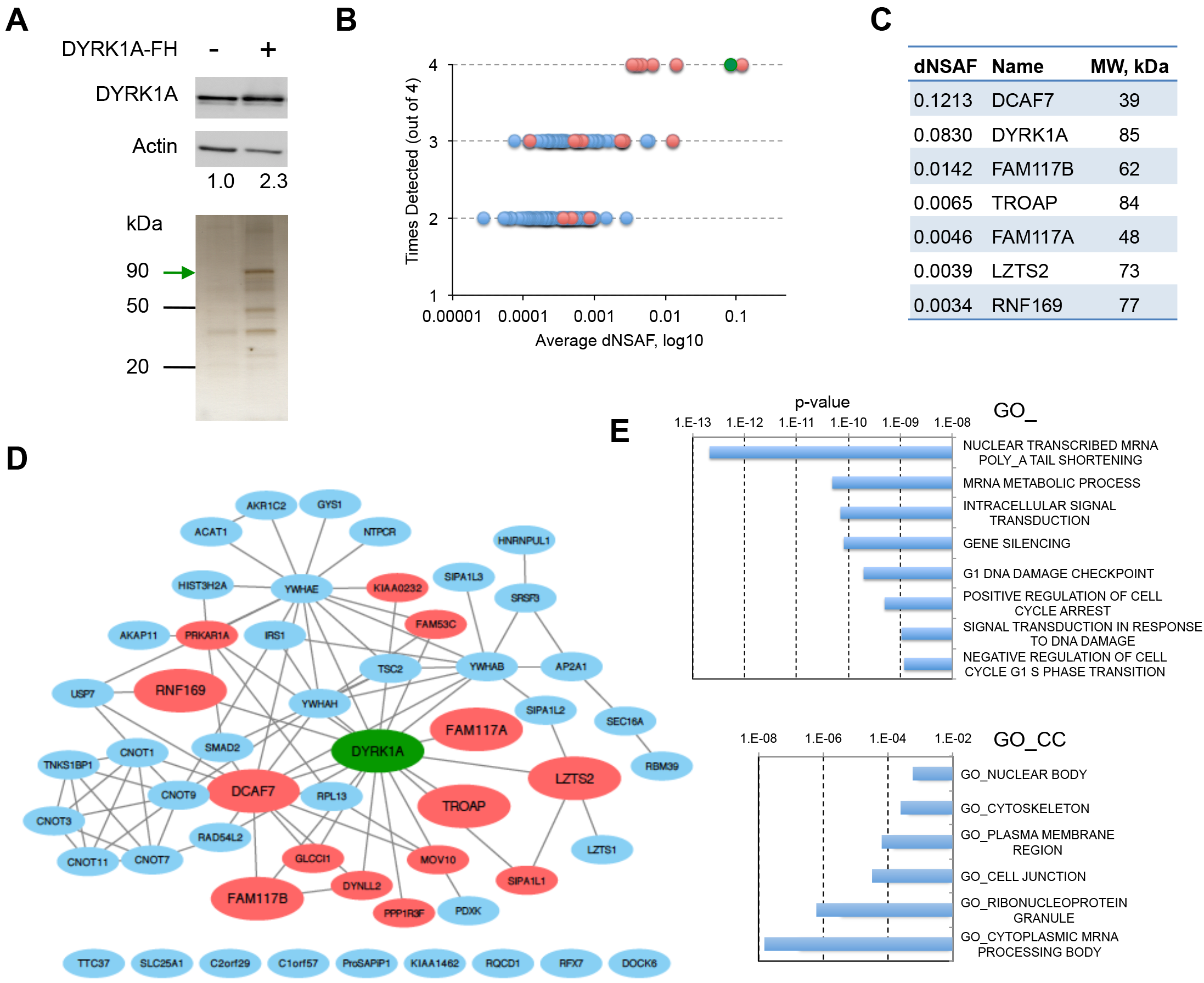
Analysis of the DYRK1A-interacting protein network. **A.** Purification of DYRK1A for MudPIT proteomic analysis. Top: Western blot showing levels of DYRK1A in T98G cells expressing HA-Flag-tagged DYRK1A (DYRK1A-FH) and DYRK1A band density relative to actin (control). Bottom: representative silver stained gel containing 10% of HA-peptide eluted control or DYRK1A-FH IP samples analyzed by MudPIT. **B.** Graph shows relative enrichment (dNSAF) of proteins detected in two, three or all four DYRK1A MudPIT experiments. DYRK1A is shown as green circle whereas red and blue circles correspond to either listed in the BioGrid database (Chatr-Aryamontri et al., 2015), or new DYRK1A-binding proteins, respectively. **C.** dNSAF (corresponds to relative enrichment) and molecular weight (MW) of seven proteins specifically detected in four DYRK1A-FH MudPIT replicate experiments. **D.** Hierarchical network of interactions (CytoScape) involving DYRK1A-binding proteins identified in this study, constructed using MetaScape analysis tool (Tripathi et al., 2015). Colors, same as in B. Larger nodes correspond to proteins detected in all four replicates. Unconnected nodes are not known to interact with other factors. **E.** Molecular Signature Database (MSigDB) annotation of the genes encoding DYRK1A-interacting proteins reveals significantly enriched functional gene ontology (GO) categories. Proteins detected in at least 3 DYRK1A MudPIT repeats were analyzed using Molecular Signature Database annotation tool to compute overlaps with GO Biological Process (GO_BP) and GO Cellular Component (GO_CC) gene sets (Subramanian et al., 2005).

### DYRK1A interacts with RNF169 and regulates recruitment of 53BP1 to DSBs

We confirmed the interaction between RNF169 and DYRK1A at both overexpressed and the endogenous levels in a series of immunoprecipitation/Western blot (IP/WB) assays using two different human cell lines, T98G and U-2 OS (Figure 2A-C). Furthermore, we found that DYRK1A and RNF169 co-fractionated together in T98G cell extract subjected to a glycerol gradient ultracentrifugation (Figure 2D). Interaction between RNF169 and DYRK1A was independent of DYRK1A’s kinase activity, and was readily detectable both in the intact U-2 OS cells and after DNA damage caused by γ-irradiation (Figure 2D). U-2 OS cells have been previously used to characterize the role of RNF169 in limiting the recruitment of 53BP1 into γ-irradiation-induced foci (IRIFs). In the preliminary time- and dose-response experiments, we found that U-2 OS cells contained the maximum number of distinct 53BP1 IRIFs at 3h after γ-irradiation (5 Gy, data not shown). Therefore, we used these conditions to assess the involvement of DYRK1A in the regulation of 53BP1 upon DNA damage in the U-2 OS cell lines stably expressing either wild type or kinase inactive DYRK1A-K188R mutant under control of doxycycline (dox)-inducible promoter (Himpel et al., 2001; Litovchick et al., 2011). As shown in Figures 2F and S2, induced expression of active, but not the kinase-inactive DYRK1A in U-2 OS cells resulted in significantly decreased number of cells displaying more than ten 53BP1 irradiation-induced foci (IRIFs) compared to un-induced control. To find out whether this effect required RNF169, we knocked down its expression in DYRK1A-inducible cell lines using siRNA (Figure 2G). Indeed, the recruitment of 53BP1 into IRIFs in the active DYRK1A-overexpressing cells was rescued to control cell levels when RNF169 expression was decreased (Figure 2H).

**Figure 2.**
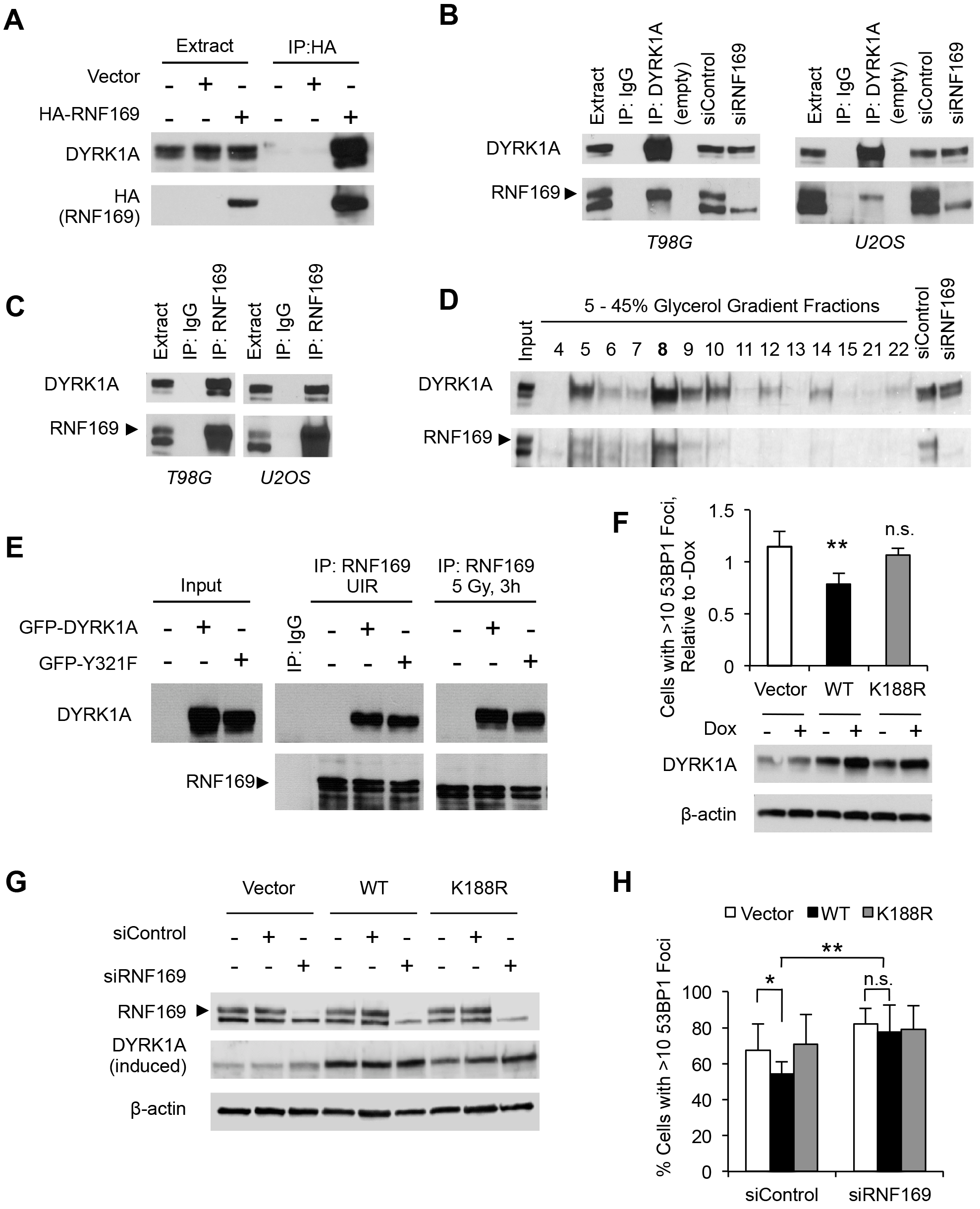
DYRK1A and RNF169 interact at the endogenous level and regulate 53BP1. **A.** IP/WB assay shows binding between transiently expressed HA-tagged RNF169 and endogenous DYRK1A in T98G cells. **B, C.** IP/WB analyses of the interaction between the endogenous RNF169 and DYRK1A in T98G and U-2 OS cells. RNF169-depleted cell extract (siRNF169) is included to identify the RNF169-specific protein band. IgG, negative control. **D.** WB of the U-2 OS cell extract separated by 5-45% glycerol gradient ultracentrifugation shows co-fractionation of RNF169 and DYRK1A. **E.** IP/WB shows that endogenous RNF169 co-precipitates the wild type and the catalytically inactive DYRK1A mutant (Y321F), from un-irradiated cells (UIR) or after DNA damage (5 Gy, 3h). **F.** Induced expression of active but not kinase-inactive K188R mutant DYRK1A inhibits the formation of 53BP1 radiation-induced foci. Inducible U-2 OS cell lines were pre-incubated with or without 1 μg/ml doxycyclin (Dox) for 12 h and then treated with radiation (5 Gy). The cells were processed for 53BP1 staining 3 h after irradiation. Graph shows quantification of % of cells with 53BP1 foci in different cell lines relative to un-induced controls (average ± stdev, N=3, Student’s two-tailed t-test results throughout the paper: n.s. or not marked − p >0.05, * − p < 0.05 and ** − p < 0.01). WB insert below the graph shows the DYRK1A levels before and after induction. **G, H.** Knockdown of RNF169 rescues DYRK1A-mediated inhibition of 53BP1 foci formation. Inducible U-2 OS cell lines were transfected with non-targeting (siControl) or RNF169-specific siRNA and treated with doxycycline to induce expression of DYRK1A. WB in panel G shows expression of proteins of interest. The graph in panel H shows quantification of the 53BP1 foci in cells collected 3h after irradiation (average ± stdev, N=4, * − p < 0.05, ** − p < 0.01).

To further investigate the role of DYRK1A kinase activity in regulating RNF169 and 53BP1, we pretreated U-2 OS parental or stably expressing HA-RNF169 cells with DYRK1A inhibitor harmine for 16 hours prior to γ-radiation, and quantified formation of the 53BP1 IRIFs. For accurate assessment, we quantified both the percentage of cells containing more than ten IRIFs, as well as an average number of IRIFs per nucleus. As seen in Figure 3A, pre-treatment of U-2 OS cells with harmine (10 μM) significantly increased the formation of 53BP1 IRIFs, supporting the role of DYRK1A in regulation of 53BP1 DSB recruitment at the endogenous levels. In agreement with previous reports, U-2 OS cells stably expressing HA-RNF169 displayed approximately 50% less cells with more than ten 53BP1 foci, and had fewer IRIFs per nucleus (Figures 3A and B, compare white bars). Treatment of these cells with harmine resulted in significant increase of 53BP1 foci formation although it was not rescued to the control levels (Figures 3A and B, compare black bars). Interestingly, HA-RNF169 foci formation was decreased by approximately 30% in harmine-treated cells compared to controls (Figures 3C and D). Since the binding between RNF169 and DYRK1A was unaffected by harmine (Figure 3E), it is likely that increased formation of 53BP1 IRIFs upon harmine treatment is due to a reduced recruitment of HA-RNF169 to DSBs in the DYRK1A-inhibited cells. Together, these results show that RNF169 and DYRK1A interact at the endogenous levels in human cells, and that DYRK1A is a positive regulator of RNF169 that limits the accumulation of 53BP1 at the DSB sites.

**Figure 3.**
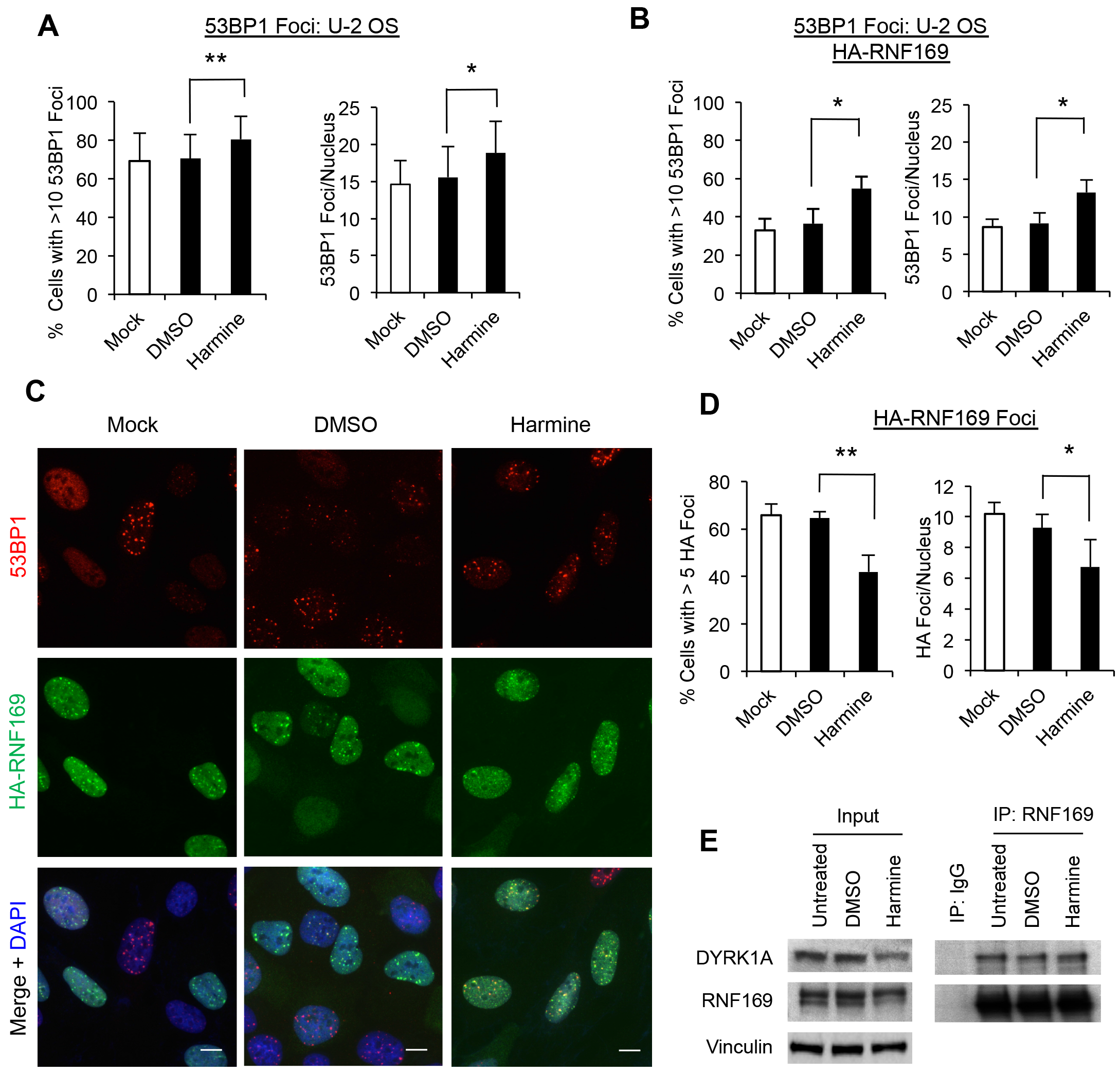
Inhibition of DYRK1A increases 53BP1 recruitment to IRIFs. **A.** Graphs show quantification of the 53BP1 radiation-induced foci in U-2 OS cells that were either untreated (Mock), or were pre-treated with 10 μM harmine or DMSO (vehicle) for 16 h before irradiation (5 Gy) and processed for staining after 3 h. Data from three experiments is shown (average ± stdev). **B.** Same as in panel A, only using U-2 OS cells stably expressing HA-RNF169.**C.** Representative images of HA-RNF169 and 53BP1 IRIFs from the experiment described in panel B. Scale bar, 10 μm. **D.** Quantification of HA-RNF169 foci in the experiment described above (average ± stdev, N=3). **E.** IP/WB analysis shows that harmine has no effect on the interaction between DYRK1A and RNF169 in U-2 OS cells.

### DYRK1A phosphorylates functionally important Ser368 and Ser403 residues in RNF169

The RNF169 protein sequence contains two predicted DYRK1A consensus sites R-x(xx)-S-P (Himpel et al., 2000). These sites, S386 and S403, are located within a highly conserved amino acid region in RNF169 that has no known function (Figure 4A). Importantly, *in vivo* phosphorylation of these sites in the native human RNF169 is reported in the PhosphoSitePlus proteomics database (Hornbeck et al., 2012). To determine whether DYRK1A can phosphorylate S368 or S403 in RNF169, we performed *in vitro* kinase assays using HA-tagged RNF169, S386A- or S403A- or S368A/S403A-RNF169 transiently expressed in HEK293T cells and immunoprecipitated, as substrates for recombinant purified DYRK1A. To avoid any co-precipitating kinase activity, HA-RNF169 immunoprecipitates were heat-inactivated at 65°C for 10 minutes prior to incubation with DYRK1A. As predicted, we observed that RNF169 was phosphorylated by DYRK1A, and that each of the S368A or S403A mutations decreased the RNF169 phosphorylation while the mutation of both sites further reduced the RNF169 phosphorylation (Figure 4B and C). It should be noted that some residual phosphorylation could be detected in the S368A/S403A (RNF169-AA) mutant. This could be due to a presence of additional, non-canonical DYRK1A phosphorylation site(s) in RNF169, or because of the contaminating kinase activity in our recombinant DYRK1A preparation.

**Figure 4.**
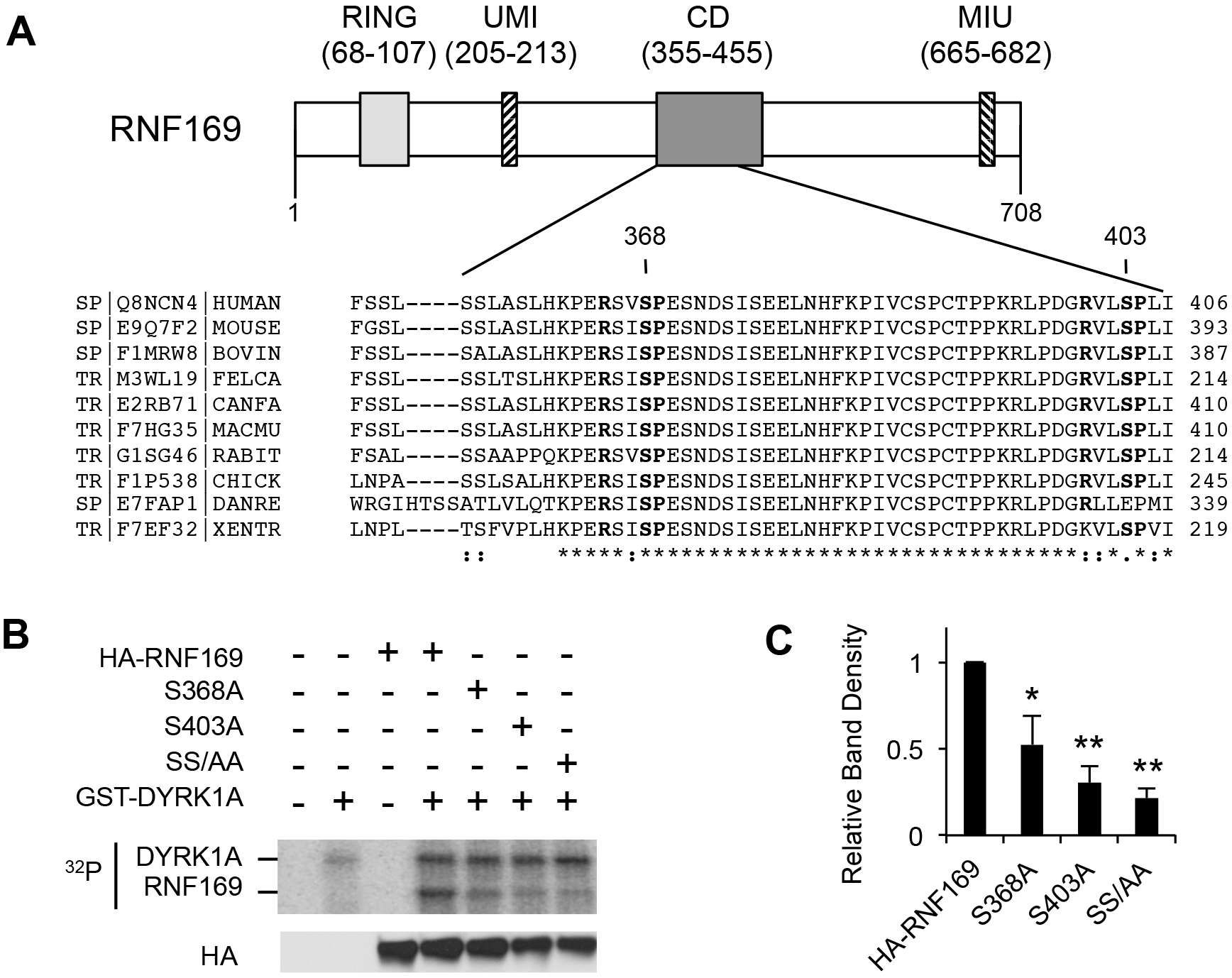
Mapping of DYRK1A phosphorylation sites in RNF169. **A.** Relative position (top) and amino acid sequence of a conserved domain (CD) in RNF169 harboring DYRK1A consensus sites (shown in bold). Amino acid numbers as in human RNF169. **B.** *In vitro* kinase assay shows that DYRK1A phosphorylates RNF169 at residues S368 and S403. HA-RNF169 or the indicated mutant proteins were expressed in HEK293Tcells, immunoprecipitated and incubated with or without purified GST-DYRK1A in a kinase reaction containing ^32^P-ATP. Proteins were resolved using SDS-PAGE and detected by phospho-imager. Bottom panel shows an aliquot of each sample analyzed by WB. Graph shows quantification of the mutant phospho-RNF169 band density relative to wild type (average ± stdev, N=3).

To further characterize the functional significance of DYRK1A phosphorylation sites in RNF169, we analyzed localization of 53BP1 and RNF169 after γ-radiation-induced DNA damage in U-2 OS cell lines stably expressing either wild type HA-RNF169, or non-phosphorylatable S368A/S340A (RNF169-AA) mutant. Interestingly, we observed approximately two-fold higher number of cells with more than ten 53BP1 foci in RNF169-AA-expressing cells compared to the wild type HA-RNF169 cell line (Figures 5A and B). The number of 53BP1 foci per nucleus was also significantly higher in the cells expressing RNF169-AA than in the wild type HA-RNF169 cells, indicating that non-phosphorylatable RNF169 mutant inhibits accumulation of 53BP1 at DSB sites to a significantly lesser extent than the wild type RNF169 (Figure 5C). Of note, the 53BP1 IRIF formation was only partially rescued in the RNF169-AA U-2 OS cell line and did not reach the control levels, similar to the effect observed in harmine-treated HA-RNF169 overexpressing cells (Figure 3B). However, while harmine treatment decreased the HA-RNF169 IRIF formation (Figure 3D), the RNF169-AA mutant showed only a slightly reduced recruitment to DSBs compared to the wild type RNF169 (Figure 5D and E). We also tested whether a phospho-mimetic mutation of the S368 and S403 to aspartic acid residues in RNF169 can rescue the 53BP1 IRIF inhibition. However, the phenotype of RNF169-DD mutant was very similar to RNF169-AA (Figure S3A, B and C). It is possible that S368D/S403D mutation does not accurately represent a constitutively phosphorylated state of the protein but instead disrupts the same function as in case of the S368A/S403A mutation. Next, we tested whether any known functions of RNF169 are impacted by mutation of the DYRK1A phosphorylation sites. Previous studies found that the recruitment of RNF169 at the DSB sites and displacement of 53BP1require ubiquitin-binding MIU domain of RNF169 that recognizes RNF168-polyubiquitilated chromatin at the site of damage (Chen et al., 2012; Hu et al., 2017; Poulsen et al., 2012). Both the RNF169-AA and RNF169-DD mutants were able to bind polyubiquitin chains similar to the wild type RNF169, further supporting our conclusion that S368 and S403 sites do not play a role in RNF169’s accumulation at the DSB sites (Figure S3E). Furthermore, mutations of these sites did not affect the interaction with ubiquitin-specific protease USP7 that has been shown to be important for RNF169 function in DNA repair (An et al., 2017) (Figure S3F). Therefore, our data presented above support the conclusion that the intact S368 and S403 sites in RNF169, together with the DYRK1A kinase activity, are required for RNF169 to fully exert its ability to limit the recruitment of 53BP1 at the DSB sites after γ-radiation, through a mechanism unrelated to recognition of the ubiquitylated chromatin or USP7 binding by RNF169.

**Figure 5.**
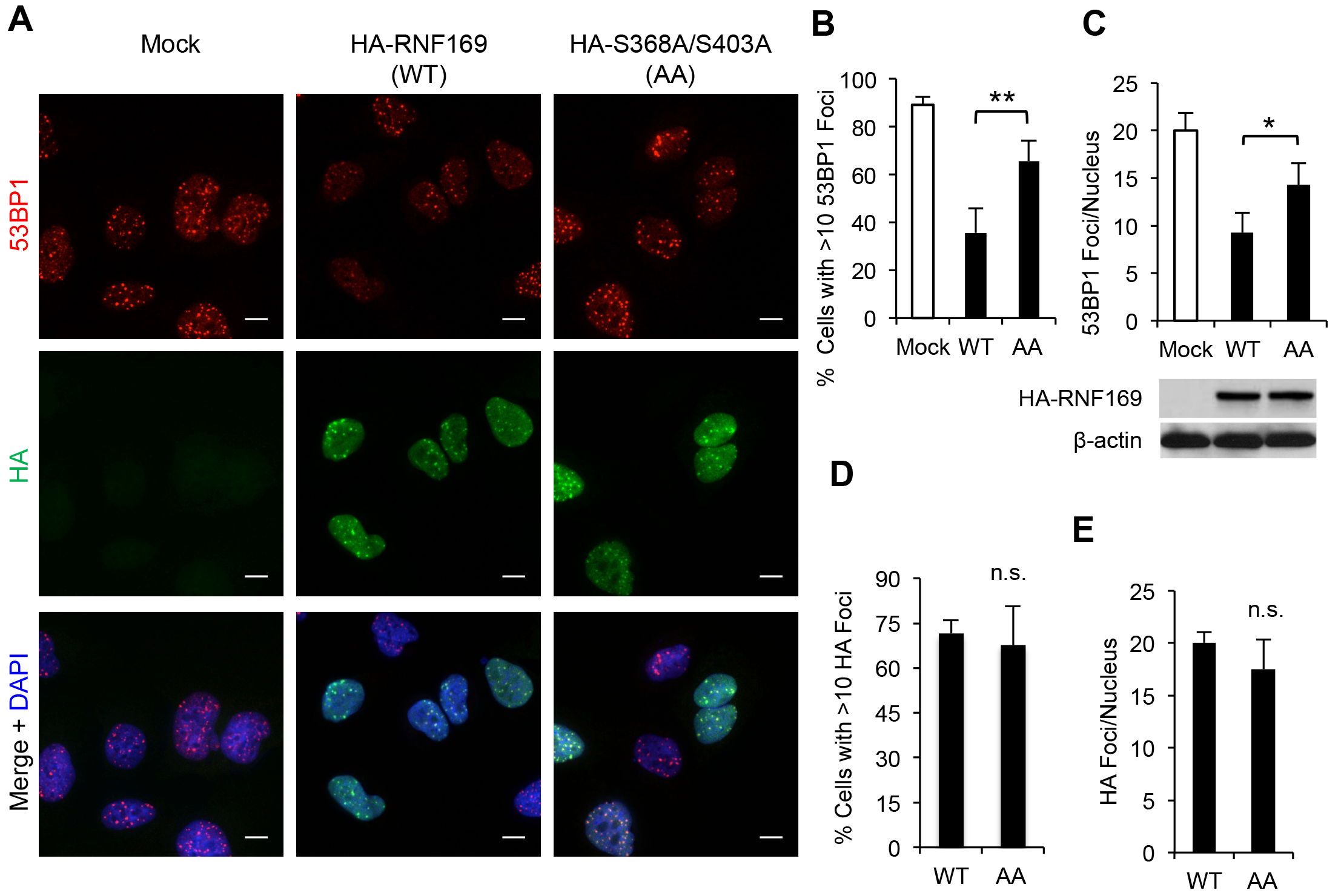
The S368 and S403 residues in RNF169 are functionally significant. **A.** Representative images of the HA-RNF169 (green) and 53BP1 (red) foci 3h after Y-radiation (5 Gy). U-2 OS stable cell lines expressing HA-RNF169 (WT) or the S368A/S403A phosphosite mutant (AA) were stained using anti-HA or 53BP1 antibodies, and DAPI. Scale bar, 10 μm. **B, C, D, E.** Graphs show quantification of the 53BP1 (B and C) and HA-RNF169 (D and E) IRIF from experiments described in A (average ± stdev, N=3). All cells were scored in the control U-2 OS cells (Mock, shown as reference) whereas only HA-positive cells were scored in the HA-RNF169 or AA-RNF169 expressing cell lines. WB under panel C shows equal expression of the WT- and AA-RNF169 constructs. Note that differences between the AA-RNF169 and wild-type RNF169 in panel D did not reach statistical significance.

### Loss of DYRK1A impairs the DSB recruitment of 53BP1 independent of RNF169

To further investigate the effects of DYRK1A loss on 53BP1 and RNF169 function, we depleted DYRK1A in U-2 OS cells using shRNA, and analyzed recruitment of HA-RNF169 and 53BP1 to DSBs. Consistent with the phenotypes observed upon inhibition of DYRK1A or mutation of the DYRK1A phosphorylation sites in RNF169, we observed an approximately 10% decrease of the HA-RNF169 IRIF formation in shDYRK1A-U-2 OS cells compared to controls (Figure S4A). However, the recruitment of 53BP1 in DYRK1A-depleted U-2 OS cells was also significantly decreased when compared to controls (Figure S4B). Furthermore, transient knockdown of DYRK1A in U-2 OS cells using two different siRNA oligos did not significantly affect the 53BP1 DSB recruitment (data not shown). Since this apparent discrepancy could be due to a residual DYRK1A expression in the siRNA or shRNA-treated cells, we generated U-2 OS cell lines harboring frame-shift mutations in *DYRK1A* gene by CRISPR-Cas9 gene editing approach (Cong et al., 2013). Of note, the Cell Line Encyclopedia data show a partial loss of chromosome 21 harboring the *DYRK1A* gene in U-2 OS (Barretina et al., 2012). Using DNA FISH and immunoblotting, we confirmed presence of a single copy of the *DYRK1A* gene in U-2 OS cells (Figure S4A). To reduce the possibility of any off-target effects, we transiently expressed Cas9 endonuclease and DYRK1A-specific guiding RNA in U-2 OS cells and isolated individual single-cell clones that were screened for loss of DYRK1A protein expression using WB. Two independent U-2 OS DYRK1A-KO clones were expanded and validated by WB and genomic DNA sequencing (Figures 6A and S5). Furthermore, we confirmed a significant loss of DYRK1A kinase activity in the U-2 OS DYRK1A-KO cell lines using a whole cell extract *in vitro* kinase assay with purified LIN52 as substrate, and a phosphospecific antibody against LIN52-S28 site for detection (Figure 6B).

**Figure 6.**
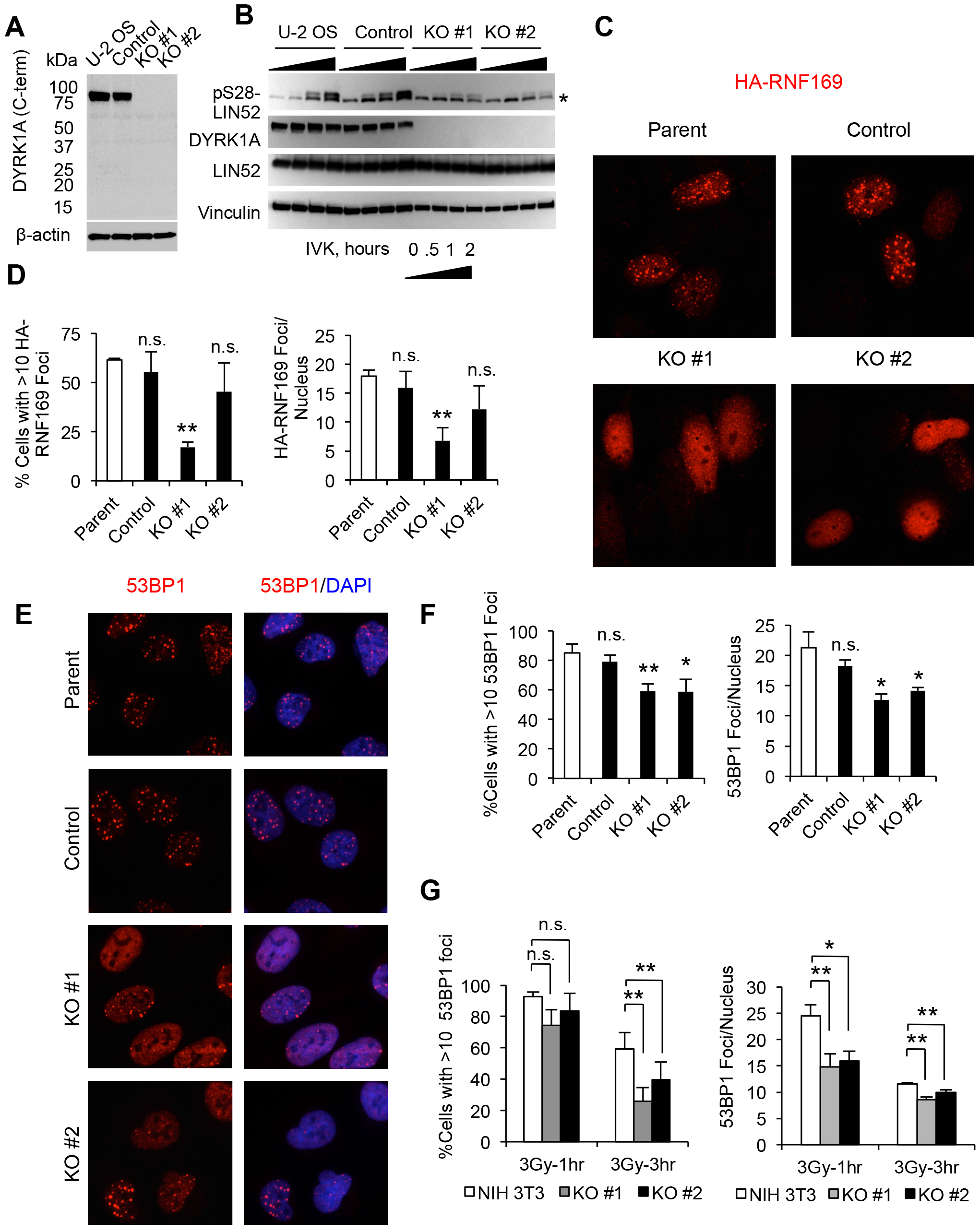
DYRK1A-deficient cells have impaired recruitment of 53BP1 to IRIFs. **A.** WB confirms absence of the full-length DYRK1A protein expression in two different U-2 OS DYRK1A-KO clones obtained using CRISPR-Cas9 approach. **B.** *In vitro* kinase assays using whole cell extracts of the control and DYRK1A-KO U-2 OS cells, and recombinant purified LIN52 as a substrate, confirms loss of DYRK1A kinase activity in the knockout cells. Phosphorylation was detected using phospho-S28-LIN52-specific antibody as described in (Litovchick et al., 2011). **C-F.** CRISPR-Cas9-mediated knockout of DYRK1A in U-2 OS cells perturbs the recruitment of 53BP1 and RNF169 to DSB sites. Representative images (C, E) and quantification results (D, F) of the 53BP1 and HA-RNF169 radiation-induced foci detected in the parental, control (non-targeting sgRNA-treated clone) as well as two different DYRK1A-KO clones. All cells were processed for staining 3h after γ-radiation (5 Gy). For statistical analysis, each cell line was compared to parental U-2 OS line (white bars) in three independent experiments. **G.** Graphs (average ± stdev, N=3) show quantification of the 53BP1 foci in two independent CRISPR-Cas9 Dyrk1a-KO clones compared to parental NIH 3T3 cells (white bars). Cells were analyzed after 1 h and 3h post γ-radiation (3 Gy).

In agreement with our observation using shDYRK1A cells, both U-2 OS DYRK1A-KO cell lines displayed a decrease in DSB recruitment of HA-RNF169 (transiently expressed) although the extent of this effect varied between the two DYRK1A-KO clones (Figure 6C and D). Furthermore, we observed a 50% decrease in the number of 53BP1 IRIFs per nucleus, resulting in significantly fewer cells with more than ten 53BP1 foci when compared to control cell lines (Fig. 6E and F). To address the mechanism of this 53BP1 recruitment defect, we analyzed the expression of several damage response markers during DNA repair. As shown in Figure S6A, there was no change in the induction of p53 or γH2AX in the DYRK1A-KO cells compared to control. Interestingly, protein levels of 53BP1 and BRCA1 appeared to be increased in the DYRK1A-KO cells compared to control whereas the expression of RNF169 was unchanged (Figure S6A). Furthermore, loss of DYRK1A in U-2 OS cells did not affect the DNA damage checkpoint, as evident by accumulation of cells in G1 and G2 phases after γ-radiation (Figure S6B). Accumulation of γH2AX and ubiquitylation of histones at the DNA damage sites also appeared to be unchanged in the DYRK1A-KO cells compared to controls (Figure S7A and B). Since accumulation of both 53BP1 and BRCA1 at the DSB sites requires the activity of RNF168 and RNF8 E3 ubiquitin ligases, we also analyzed the recruitment of BRCA1 in these cells and found it similar (Figure S7C). Therefore, we concluded that decreased recruitment of 53BP1 to the damage sites was likely not because of abnormal DNA damage signaling or histone ubiquitylation in the DYRK1A-KO cells.

Since U-2 OS cells could have intrinsically reduced DYRK1A expression resulting from loss of one copy of the *DYRK1A* gene, we sought to further validate our findings described above using a different cell line. To do so, we knocked out Dyrk1a expression in mouse NIH-3T3 fibroblasts using CRISPR-Cas9 approach (Figure S5D). We first determined that the number of 53BP1 foci in NIH-3T3 cell peaked after 1h post γ-irradiation and started to decrease at 3h, and that 3 Gy was the optimal dose to observe distinct nuclear foci (data not shown). Using these experimental conditions, we again observed an impaired 53BP1 IRIF formation in two independent single-cell derived NIH-3T3 Dyrk1a-KO clones compared to control cells (Figure 6G). Interestingly, while approximately 80% of both the control and Dyrk1a-KO NIH-3T3 cells contained more than 10 53BP1 foci at 1h post Y-irradiation, initial recruitment of 53BP1 was impaired by loss of Dyrk1 a because the average number of foci per nucleus was significantly lower in the Dyrk1a-KO cells compared to control (Figure 6G). At 3h post γ-irradiation, the number of foci per nucleus decreased by approximately 50% in both the control and Dyrk1a-KO cell lines, indicative of a similar rate of resolving the lesions and removal of 53BP1 foci in these cells lines.

To further confirm that the phenotype of the DYRK1A-KO U-2 OS cells was specific to a loss of DYRK1A protein and kinase activity, we re-introduced either active DYRK1A, or kinase inactive K188R-DYRK1A mutant, into one of our knockout clones. Indeed, we observed that expression of the wild-type DYRK1A, but not the kinase inactive mutant, resulted in a complete rescue of the 53BP1 IRIF defect in these cells (Figures 7A and B). We also investigated whether the effect of DYRK1A loss on 53BP1 is mediated by RNF169. Interestingly, unlike DYRK1A overexpressing cells (Figure 2H), knockdown of RNF169 in DYRK1A-KO U-2 OS cells resulted in only a partial rescue of the 53BP1 recruitment to the DSB sites, and the RNF169-depleted DYRK1A-KO cells still showed significantly lower 53BP1 IRIF formation when compared to corresponding control cell lines (Figure 7D). This result suggests that the 53BP1 recruitment defect in the absence of DYRK1A is not likely due to a more efficient displacement by RNF169. Together, these results support the conclusion that DYRK1A regulates the initial recruitment of 53BP1 to damaged chromatin both in the RNF169-dependent and independent manner.

**Figure 7.**
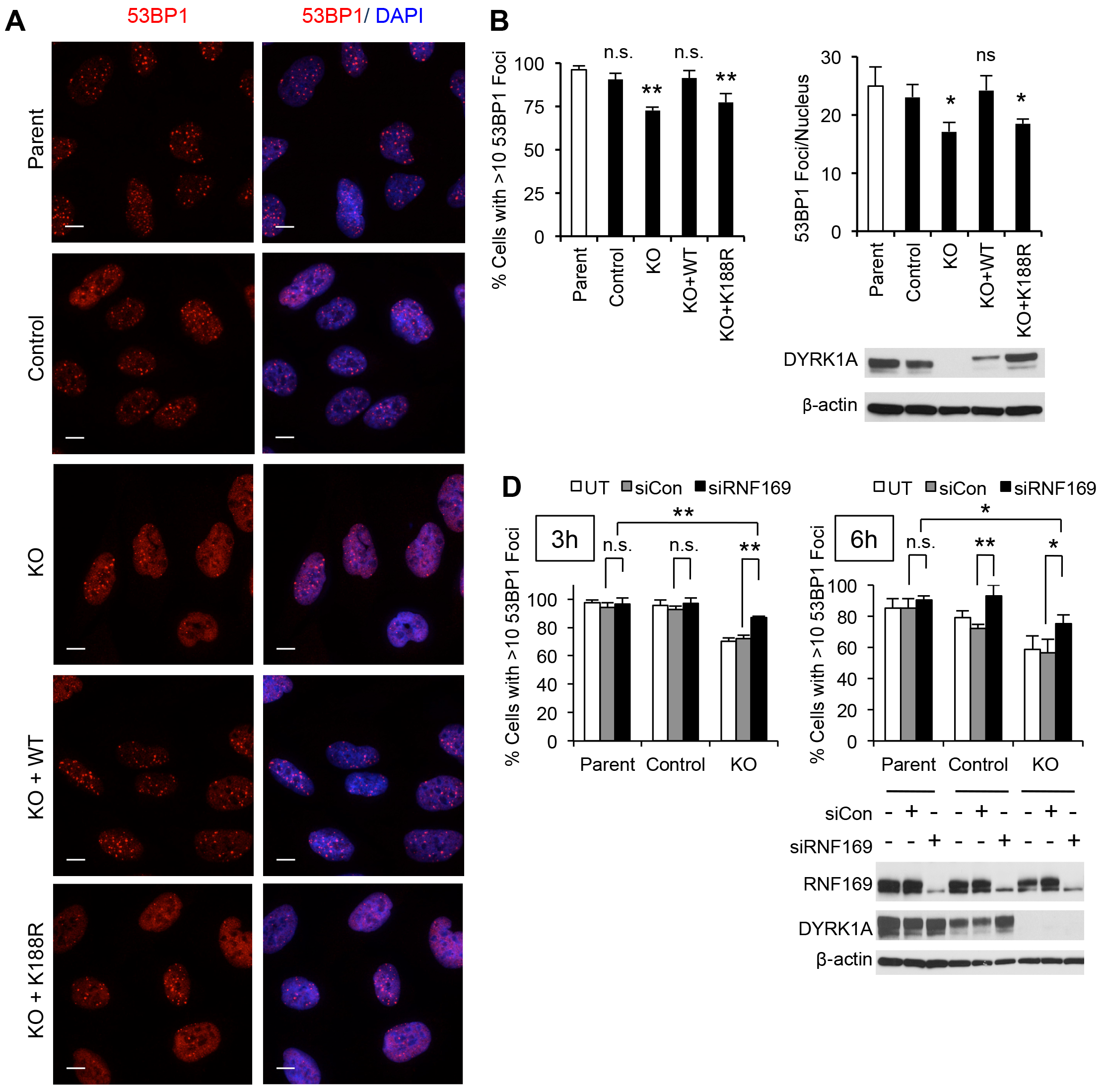
Rescue of the 53BP1 foci formation in DYRK1A-KO cells by re-expression of active DYRK1A or depletion of RNF169. **A.** Representative images of 53BP1 staining in parental U-2 OS cells, control sgRNA clone, in DYRK1A-KO clone #1 (KO), as well as KO clone after stable re-expression of the wild type (KO + WT) or kinase-inactive mutant DYRK1A (KO + K188R). Scale bar, 10 μm. **B, C.** Graphs show quantification (average ± stdev) of the 53BP1 radiation-induced foci from three independent experiments using the cell lines described in A. For statistical analysis, each cell line was compared to parental U-2 OS line (white bars). WB in panel C confirms re-expression of DYRK1A in the KO cells. Slower gel migration of the recombinant DYRK1A alleles is due to the presence of dual Flag-HA tag. **D.** Graphs show quantification (average ± stdev, N=3) of the 53BP1 radiation-induced foci in control or DYRK1A-depleted U-2 OS cell lines after siRNA knockdown of RNF169. For statistical analysis, siRNF169-treated cells were compared to non-targeting siRNA-transfected cells (siControl). Un-transfected cells (UT) are shown as reference (white bars). WB in panel E shows the expression levels of RNF169 and DYRK1A in the representative experiment.

### Loss of DYRK1A promotes the HRR and DNA repair

Previous studies demonstrated the role of 53BP1 in suppressing the HR-mediated DNA repair by protecting the DNA ends around the site of damage from resection (Canny et al., 2018; Durocher and Pelletier, 2016). Since loss of DYRK1A decreased accumulation of 53BP1 at the DSB sites, we investigated its effect on the DNA repair pathway determination using the control and DYRK1A-KO U-2 OS cell lines stably expressing the direct repeat (DR) GFP reporter of the HR repair (Pierce et al., 1999). In this model system, a DSB is generated by cleavage of the non-functional GFP gene fragment by I-SceI restriction nuclease. The break is then repaired either by NHEJ, resulting in no GFP protein expression, or by HRR, in which case a fluorescent protein is produced (Fig. 8A). Consistent with reduced recruitment of 53BP1 to DSBs in the DYRK1A-KO cells, we observed approximately two fold increase in GFP-positive cells formation after I-SceI expression in these cells compared to control U-2 OS cells (Fig. 8B).

**Figure 8.**
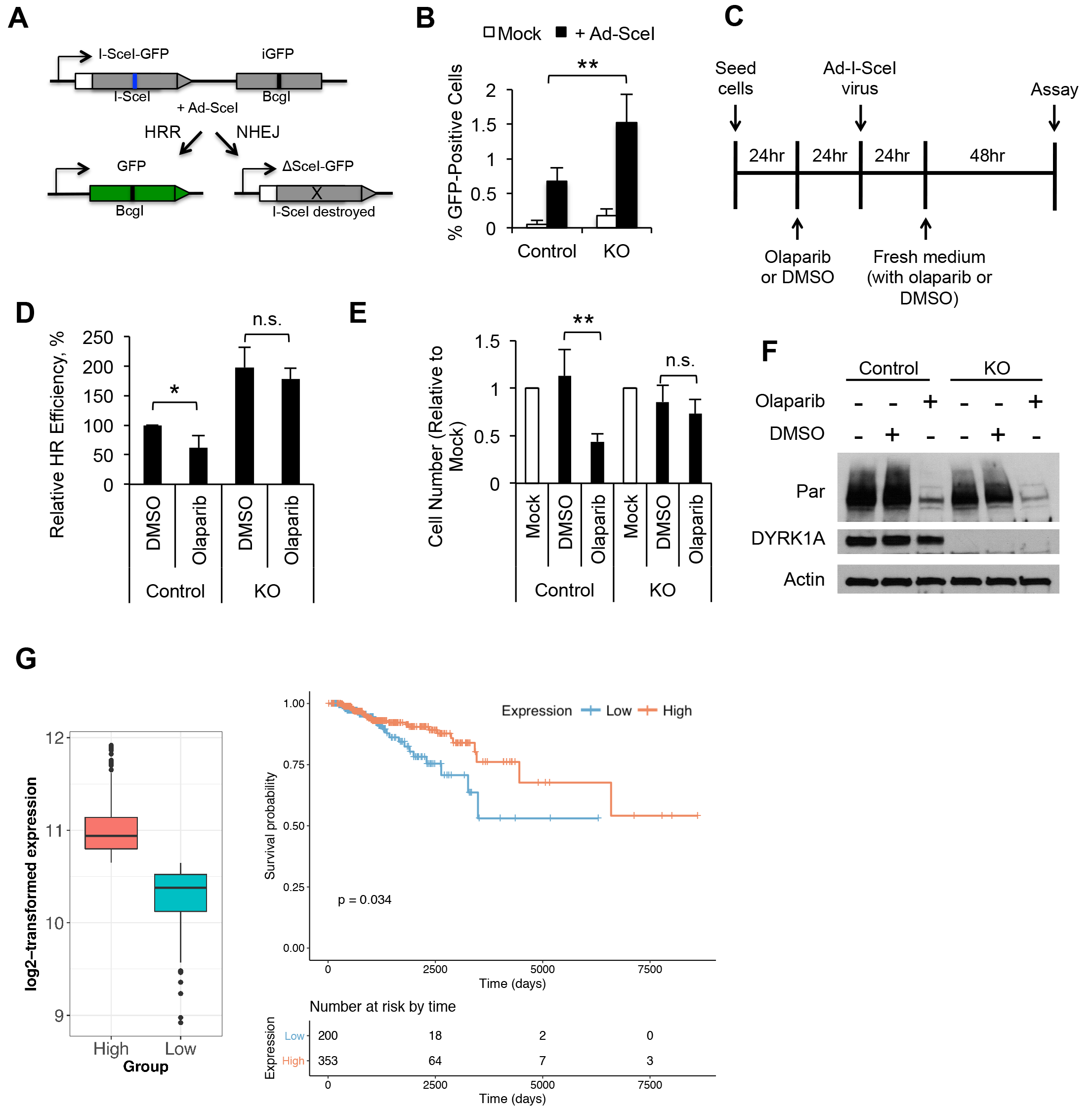
Loss of DYRK1A promotes the HRR and increases resistance to DNA damaging therapy. **A.** Schema of the DR-GFP reporter that produces fluorescent protein only after it is cut by I-SceI endonuclease and repaired by HRR (Pierce et al., 1999). **B.** U-2 OS DYRK1A-KO cells (clone #1) display an increased HRR repair efficiency compared to control U-2 OS cells as measured by DR-GFP reporter expression. DR-GFP reporter was stably expressed in the control or DYRK1A-KO cells using transfection followed by antibiotic selection. I-SceI was introduced using adenovirus infection. Graph shows the results of FACS analysis measuring the percentage of GFP-positive cells in the total population (average± stdev, N=3). **C.** Schema of the cell treatments for the experiments shown in panels D, E and F. **D.** Olaparib treatment (8 μM) reduces the HR repair efficiency in the control but not in the DYRK1A-KO U-2 OS cells. HR repair efficiency was determined using DR-GFP reporter cell lines as described in panel B, and the % of the GFP-positive cells was normalized to that of control cells treated with DMSO (vehicle). For statistical analysis, olaparib-treated cells were compared to DMSO-treated cells in three independent experiments. **E.** DYRK1A-KO cells display greater resistance to growth inhibitory effect of olaparib when compared to control cells. Graph shows changes in cell numbers (normalized to untreated cells, Mock) after incubation with either DMSO or 8 μM olaparib. **F.** WB confirms efficient inhibition of PARP activity in the control and DYRK1A-KO cells under the experimental conditions shown in panels D-F. **G.** Low expression of DYRK1A in breast cancer is associated with worse survival in patients treated with radiotherapy. Boxplot graph shows DYRK1A mRNA levels in the highest 25% (High) and the lowest 25% (Low) expression groups of the TCGA breast cancer samples. Kaplan-Meier survival plot shows that patients from “Low” DYRK1A group had significantly worse overall survival than “High” DYRK1A group.

Targeting DNA repair processes is an important strategy for anti-cancer therapy (Brown et al., 2017). Extensive studies established that HR-mediated repair is the principal mechanism of DNA repair in cells treated with inhibitors of anti-cancer drugs inhibiting poly (ADP-ribose) polymerases (PARP), such as olaparib, making these drugs especially efficient in cancers with mutations in the HR repair pathway (Lin and Kraus, 2017; Pommier et al., 2016). In HR-proficient cancer cells, olaparib can potentiate the effect of other DNA damaging drugs (Griguolo et al., 2018; McCann, 2018), possibly by attenuating the efficiency of HR-mediated repair as it was shown in U-2 OS cells (Jelinic and Levine, 2014). Factors that mediate cancer cell resistance to PARP inhibitors are not fully understood; therefore, we were interested to see if loss of DYRK1A can influence the cell response to olaparib by promoting HR (Figure 8C). Consistent with previous data, in case of control DR-GFP U-2 OS cells, olaparib caused a significant inhibition of the HR-mediated repair efficiency and decrease of cell number as compared to vehicle-treated cells. However, DYRK1A-KO cells appeared to be more refractory to olaparib effect than the control cells although the extent of PARP inhibition was similar in both cell lines (Figure 8C-F). Therefore, loss of DYRK1A in cancer cells could increase resistance to PARP inhibitors or other DNA damaging treatments by facilitating the HR-mediated repair.

In order to determine whether different DYRK1A mRNA expression levels in cancer can be associated with particular clinical outcomes, we searched publically available cancer TCGA datasets. We found that expression of DYRK1A was significantly reduced in cancer samples compared to normal tissue in several cancer types, including invasive breast carcinoma (data not shown). Interestingly, while DYRK1A mRNA expression had no significant association with the outcomes in the entire breast cancer patient’s dataset, low expression of DYRK1A significantly correlated with decreased survival in the group of patients treated with γ-irradiation (Figure 8G). Similar results were observed in radiation-treated lung adenocarcinoma patient cohort (data not shown), indicating that low DYRK1A expression could be a negative prognostic marker for predicting the outcomes of radiation therapy in certain types of cancer.

## Discussion

*DYRK1A* gene copy number changes have deleterious effects on prenatal and early postnatal brain development and have been linked to neurodegenerative disease and cancer (Abbassi et al., 2015; Dierssen and de Lagran, 2006; Tejedor and Hammerle, 2011). Since even subtle changes in DYRK1A levels appear to deregulate its function, it is possible that some effects of *DYRK1A* gene imbalance could be mediated by perturbation of DYRK1A interaction networks. Our data presented here offer an insight into a complexity and functional diversity of the protein-protein interactions that involve this remarkable protein kinase in human cells.

Currently, the BioGrid protein interaction network database lists 80 DYRK1A-interacting proteins, mostly identified by high-throughput affinity-capture mass spectrometric analyses performed in HEK293T cells (Chatr-Aryamontri et al., 2015). Using sensitive MudPIT proteomic approach (Florens and Washburn, 2006; Swanson et al., 2009), we identified 120 proteins specifically detected in at least two out of four biological replicate analyses of DYRK1A immunoprecipitates from human T98G cells, including 98 proteins not reported to interact with DYRK1A in the BioGrid protein interaction database. Given evidence of the functional significance of DYRK1A in T98G cells (Di Vona et al., 2015; Litovchick et al., 2011; Litovchick et al., 2007), our new data on the DYRK1A protein-protein interaction network in human cells will serve as a resource for the future functional studies of this important protein kinase.

Our analysis identified DCAF7 (also known as WDR68 or HAN11) as most highly enriched in the DYRK1A immunoprecipitates among DYRK1A-binding proteins. Structurally, DCAF7 is a WD40-repeat protein that directly binds to several protein kinases including DYRK1A, and serves as an adaptor to mediate their interactions with other proteins including adenovirus E1A protein (Glenewinkel et al., 2016; Miyata and Nishida, 2011; Ritterhoff et al., 2010). Some of DYRK1A interactions identified in our study could be mediated by DCAF7, and further studies will be needed to characterize the role of DCAF7 in regulating the DYRK1A interaction networks. Furthermore, the DCAF7-mediated interaction of DYRK1A with E1A could alter DYRK1A interacting networks in HEK293T cells because of the presence of this viral protein. Indeed, while fourteen of the DYRK1A-interacting proteins were detected in both T98G and 293T cells (Figure S1A), several of the proteins known to interact with E1A including RB1, RBL1, RBL1 and EP300, were not detected in our analysis ((Varjosalo et al., 2013) and this work). In addition to its adaptor function, DCAF7 is recruited into the Cul4-DDB1 ubiquitin ligase complex that has been recently shown to regulate stability of DNA ligase I (LIG1), one of the key enzymes in the alternative NHEJ DNA repair (Peng et al., 2016). It remains to be determined whether DYRK1A plays a role in the DCAF7-mediated degradation of LIG1 and DNA repair. Furthermore, the proteins that are most enriched in the DYRK1A interactome remain to be characterized, and their functional connection to DYRK1A is not apparent. Also of note, DYRK1A-interacting proteins TROAP and LZTS2 have been shown to function as tumor suppressors (Cui et al., 2013; Johnson et al., 2013; Lian et al., 2017), and it will be important to determine their role in DYRK1A-mediated inhibition of cell proliferation (Litovchick et al., 2011).

The complexity of the protein-protein interaction network involving DYRK1A likely reflects its diverse functions in the context of specific cellular compartments or as part of different protein complexes. Here, we provide functional characterization of the interaction between RNF169 and DYRK1A that revealed the role of DYRK1A in the DNA damage pathway by regulating 53BP1, one of the key response factors to DNA DSB lesions (Panier and Boulton, 2014). The DNA DSBs are repaired in the cell cycle-dependent manner by either homologous recombination repair (HR) or the non-homologous end joining (NHEJ), and the choice of appropriate repair mechanism involves multiple factors that mediate and recognize modifications of the chromatin around the lesion (Hustedt and Durocher, 2016). Ubiquitin ligase RNF168 plays a key role in recruitment of 53BP1 to DSB sites (Doil et al., 2009; Fradet-Turcotte et al., 2013; Panier and Boulton, 2014). RNF168 binds to RNF8-ubiquitylated histones and catalyzes H2A-K15ub modification required for the recruitment of 53BP1 that protects the damaged DNA ends from excision and therefore facilitates the repair by NHEJ (Doil et al., 2009; Panier and Durocher, 2009). RNF169 is a homolog of RNF168 and a relatively new player in the DNA damage response pathway. RNF169 also recognizes H2A-K15ub marks but lacks the E3 ubiquitin ligase of its own, therefore its accumulation is thought to limit the recruitment of 53BP1 to DSBs (Chen et al., 2012; Hu et al., 2017; Poulsen et al., 2012). The function of RNF169 is best revealed upon its overexpression when it prevents the accumulation of 53BP1 at the DSBs, resulting in increased HR-mediated DNA repair efficiency due to a more efficient resection of the DNA ends (Chen et al., 2012; Poulsen et al., 2012). However, the mechanism of the RNF169 activity towards 53BP1 is not fully understood.

Our study confirms the role of RNF169 as a negative regulator of 53BP1 accumulation, and supports the role of DYRK1A as an RNF169 kinase that positively regulates this activity through both direct and indirect mechanisms. Previous studies found that a high-affinity ubiquitin-binding MIU domain in RNF169 is required for its ability to inhibit 53BP1 accumulation at the damage sites (Chen et al., 2012; Hu et al., 2017; Poulsen et al., 2012). Our study extends this observation by demonstrating that while the binding of RNF169 to the ubiquitylated histones surrounding the DSBs could be necessary, it is not sufficient to prevent the accumulation of 53BP1 at the DSB sites. Indeed, the phosphorylation-deficient RNF169 mutants show reduced ability to displace 53BP1 from the DSBs despite apparently normal recruitment to these sites. Interestingly, seven ATM-regulated phosphorylation sites in 53BP1 are required for its interaction with its key effector RIF1 but dispensable for its recruitment to the damage sites (Escribano-Diaz et al., 2013; Isono et al., 2017). It is possible that DYRK1A phosphorylation of RNF169 serves to recruit an additional factor that is essential for displacing 53BP1, or for stabilizing the binding of RFN169 to ubiquitylated chromatin. Constitutive presence of the DYRK1A-RNF169 complex in the intact cells and after damage also indicates that another factor could be recruited during DNA damage response. Further proteomic studies of the RNF169-DYRK1A complex in the cells before and after DNA damage will help to identify such factor. Our analysis of DYRK1A detected interaction with USP7, a ubiquitin-specific protease that has been recently shown to bind directly to RNF169 and to increase the stability of 53BP1, RNF169 and RNF168 (An et al., 2017; Liu et al., 2016; Yim et al., 2017; Zhu et al., 2015). Although disruption of the DYRK1A phosphorylation sites in RNF169 did not influence its interaction with USP7, the role of USP7 in the DYRK1A-RNF169 mediated regulation of 53BP1 should be further investigated.

Interestingly, while DYRK1A can facilitate the displacement of 53BP1 from DSBs by overexpressed RNF169, the 53BP1 DSB recruitment defect in the DYRK1A-depleted cells appears to be, at least in part, RNF169-independent. Indeed, DYRK1A-KO cell lines displayed decreased RNF169 IRIF formation, and the 53BP1 recruitment phenotype could not be fully rescued by RNF169 depletion in these cells. Recent studies revealed that in addition to histone H2A-K15ub mark, 53BP1 recognizes and binds to H4K20Me2 mark via its conserved Tudor domain, and this process is regulated by several factors including histone methyltransferases SETD8 and MMSET, as well as Polycomb proteins L3MBTL1 and JMJD2A that occupy these marks in the absence of DNA damage (reviewed in (Panier and Boulton, 2014)). Furthermore, in S/G2 phases of the cell cycle, BRCA1 plays an active role in removing 53BP1 from chromatin around the damage sites using a complex and not fully understand mechanism that requires CDK activity and CtIP (Chapman et al., 2012; Escribano-Diaz et al., 2013; Isono et al., 2017). Interestingly, BRCA1 was upregulated and efficiently recruited to DSB sites in DYRK1A KO U-2 OS cells therefore it is possible that BRCA1-mediated eviction of 53BP1 could contribute to the phenotypes observed in these cells. Since BRCA1 gene expression could be regulated by DYRK1A through recruitment of the DREAM repressor complex (Litovchick et al., 2011; Litovchick et al., 2007; Yakovlev, 2013), the relationship between DYRK1A expression levels and the outcomes of the DNA damaging therapy in cancer should be further investigated. Interestingly, loss of 53BP1 can rescue the HR defects associated with inactivation of BRCA1, and could be responsible for the acquired resistance of the BRCA1-mutant tumors to PARP-inhibitor therapy (Jaspers et al., 2013). Therefore, future studies will be needed to establish the exact role of DYRK1A in the context of cellular processes that regulate the recruitment of 53BP1 to the DSBs, and to validate the significance of DYRK1A as a factor that can influence the outcomes of cancer therapy.

## Experimental Procedures

### Cell lines

Human osteosarcoma U-2 OS, glioblastoma T98G, HEK293T and mouse NIH3T3 cells were obtained from ATCC and used from early passage master stocks. Cells were regularly checked for mycoplasma using in-house PCR assay and DAPI staining. T98G cells stably expressing Flag-HA epitope tagged DYRK1A, GFP or DYRK1A-interacting proteins were established using pMSCV retroviral vectors and puromycin selection as described in (Litovchick et al., 2011); this work also describes the doxycycline-inducible U-2 OS cell lines expressing DYRK1A. DYRK1A-null U-2 OS and NIH3T3 cells were established using GeneArt CRISPR Nuclease vector with OFP reporter (Life Technologies) harboring human or mouse DYRK1A-specific guide sequences. The control cell line was similarly established using a non-targeting construct provided with the kit. Briefly, cells were transfected with sgRNA-CRISPR plasmids, FACS-sorted for OFP expression and grown as single-cell clones that were screened for DYRK1A expression using immunoblotting. Two different clones lacking DYRK1A expression were expanded and validated using antibodies against different epitopes in DYRK1A as well as genomic sequencing of the nested PCR-amplified fragment surrounding the sgRNA-targeted region. Human DYRK1A-specific guiding sequence for CRISPR-Cas9 genomic mutagenesis: top strand: 5′-tgtaaaggcatatgatcgtg-3′ and bottom strand: 5′-cacgatcatatgcctttaca-3′. Nested primers for PCR amplification of *DYRK1A* genomic region 400 bp up- and downstream of the Cas9 targeting site: first PCR set: forward 5′-aagttatctgaagccttctgc-3′ and reverse 5′-catggtatgctacatggaaggc-3′; second PCR set: - forward 5′-cttagggttcaggtatctctc-3′ and reverse 5′-ccaagatttagactattactac-3′). The second PCR primer set was also used for sequencing of the purified PCR products. Mouse Dyrk1a-specific guiding sequence for CRISPR-Cas9 genomic mutagenesis: top strand: 5′-ggacgattccagtcataaga-3′, and bottom strand: 5′-tcttatgactggaatcgtcc-3′. Nested primers for PCR amplification of DYRK1A genomic region 480 bp up- and downstream of the Cas9 targeting site: first PCR primer set: forward 5′-gaacattgagttcaactttgaggg-3′ and reverse 5′-ggcactgactagccagaaacc-3′; second PCR primer set: forward 5′-ttgtttgggggttccttgtg-3′ and reverse 5′-caagaagtggcagcttgctg-3′. To verify clonal origin of established KO cell lines, the amplified genomic regions from the PCR with 2nd primer set were purified, cloned into Promega pGEM^®^-T Easy vector. Multiple DNA clones were sequenced to confirm the presence of mutations and the absence of the wild type sequences.

### Chemicals and treatments

To induce DNA damage, the cells were exposed to gamma irradiation using MDS Nordion Gammacell 40 research irradiator with a Cs^137^source (ON, Canada) or subjected to laser beam irradiation to induce DNA damage as described in (Poulsen et al., 2012). Olaparib (AZD2281, Ku-0059436 from Selleckchem, catalog No. S1060) and harmine (from Sigma; catalog No. H8646) stock solutions were prepared in DMSO, and used for cell treatments at 8μM or 10μM final concentrations, respectively.

### RNAi and Plasmids

siRNA oligos used in this study were from Ambion/Thermo Fisher Scientific including siRNF169 (Silencer Select, ID: s48512, Cat# 4392420) and Negative Control No.1 siRNA (Silencer Select, Cat# 4390843). siRNA transfections were performed using Lipofectamine RNAiMAX (Invitrogen) according to the manufacturer’s instructions. GFP-tagged mouseDyrk1a wild-type and mutant constructs were a kind gift from G. D’Arcangelo (Yabut et al., 2010). HA-RNF169-pcDNA3 construct was a gift from N. Mailand (Poulsen et al., 2012). The phospho site mutants of RNF169 were generated using the QuikChange II XL site directed mutagenesis kit (Agilent Technologies), and verified by sequencing. Plasmid transfections were performed using either TransIT-2020 transfection reagent (Mirus Bio) or polyethylenimine reagent (Polysciences Inc) that was prepared according to manufacturer’s protocol.

### MudPIT proteomic analysis

MudPIT proteomic analysis was performed as described in (Florens and Washburn, 2006; Litovchick et al., 2011; Litovchick et al., 2007) using Finnigan LTQ Linear ion trap mass spectrometer equipped with an electrospray ionization source. T98G cells stably expressing DYRK1A-Flag-HA or GFP-Flag-HA (control) were used for immunoprecipitations with anti-HA antibody agarose beads (clone HA7, Sigma). Proteins were eluted from beads using HA peptide, concentrated and digested with trypsin. Tryptic peptides were resolved using Quaternary Agilent 1100 series HPLC and microcapillary multi-dimensional C_18_-SCX-C_18_ matrix using fully automated 10-step chromatography run and electrosprayed into mass spectrometer. Full MS spectra were recorded on the peptides over a 400 to 1,600 m/z range, followed by five tandem mass (MS/MS) events sequentially generated in a data-dependent manner on the first to fifth most intense ions selected from the full MS spectrum (at 35% collision energy). SEQUEST(Eng et al., 1994) was used to match MS/MS spectra to peptides in a database of 58622 amino acid sequences, consisting of 29147 Human proteins (non-redundant entries from NCBI 2011-08-16 release). To estimate relative protein levels, spectral counts were normalized using Normalized Spectral Abundance Factors (NSAFs) (Litovchick et al., 2007; Swanson et al., 2009; Zybailov et al., 2006). Average NSAFs were calculated from four biological replicate DYRK1A pull-down experiments.

### Kinase assays

For RNF169 phosphorylation assays, HEK293T cells were transfected with HA-tagged RNF169 constructs and lysed using RIPA buffer (50mM Tris-HCl, pH 7.4, 150 mM NaCl, 1% NP-40, 0.5% Sodium deoxycholate and 0.1% SDS) supplemented with phosphatase and protease inhibitor cocktails (Millipore) and 1:10,000 β-mercaptoethanol. Extracts were incubated with 1 μg anti-HA antibody and protein A beads followed by six washes in EBC buffer (50 mM Tris-HCl, pH 8.0, 5 mM EDTA, 120 mM NaCl and 0.5% NP-40, phosphatase and protease inhibitors and 1:10,000 β-mercaptoethanol, and one final wash with kinase assay buffer (Cell Signaling) containing 25 mM Tris-HCl, pH 7.5, 5 mM β-glycerophosphate, 2 mM DTT, 0.1 mM Na_3_VO_4_, 10 mM MgCl_2_. The immunoprecipitated wild type or mutant RNF169 proteins were used as substrates in a kinase assay reaction in the presence of 1 μM cold ATP, 5 μCi of [γ-^32^P] ATP and 200 ng GST-DYRK1A (Life Technologies) for 30 min at RT. The beads containing phosphorylated proteins were washed once with EBC buffer and analyzed by SDS-PAGE and autoradiography. For LIN52 phosphorylation assays, extracts from control and DYRK1A-KO U-2 OS cell lines (1 mg/ml) were prepared using EDTA-free EBC buffer supplemented with phosphatase inhibitors, 2 mM DTT, 10 mM MgCl2, 10 mM MnCl2 and 200 μM ATP, and incubated at 30°C with 6 ng GST-LIN52 in a 100 μl reaction volume. Reactions were terminated at different times by adding SDS-PAGE loading buffer and heating at 95°C for 10 min, and analyzed by WB with indicated antibodies as described in (Litovchick et al., 2011).

### Immunofluorescence

Cells were seeded on glass coverslips in 6-well dishes and allowed to attach for 24 h. After washing in PBS three times, cells were fixed in 4% formaldehyde for 20 min and permeabilized with 0.2% Triton X-100 in PBS containing 5% BSA for 30 min followed by incubation with primary and secondary antibodies (1h at room temperature). The coverslips were then mounted in Fluoroshield mounting medium with DAPI (Abcam) and viewed using Zeiss Axio AX10 Imager.M1m fluorescence microscope equipped with AxioCam MRm camera. Images were acquired using AxioVision, AxioVs40 (V 4.8.2.0) software. The images were analyzed using FIJI software (Schindelin et al., 2012). Briefly, images in JPEG format were processed to find the total number of foci (maxima). A noise tolerance value of 20 or 30 was used, and it was the same for all samples within each comparison group. For 53BP1, average foci per cell and number of cells with greater than 10 foci were calculated. For HA-RNF169, average foci per HA-positive cell and number of HA-positive cells with greater than 5 or 10 foci were calculated. To analyze 53BP1 in HA-RNF169 expressing cell lines, 53BP1 foci were scored only in the HA-positive cells. For each biological repeat, more than 100 cells were typically processed.

### Immunoblotting and immunoprecipitation

For immunoblotting, cells were lysed in EBC or RIPA buffers for 10 min at 4 °C and then centrifuged at 14,000g for 15 min at 4 °C. Protein concentrations were determined by DC protein assay (BioRad). Protein samples were resolved using polyacrylamide gels (BioRad), transferred to nitrocellulose membrane (GE Healthcare) and probed by specific antibodies as recommended by manufacturer. For immunoprecipitation, cell extracts were incubated with appropriate antibodies (1 μg/ml) and Protein A Sepharose beads (GE Healthcare) overnight at 4 °C, washed five times with lysis buffer and re-suspended in Laemmli sample buffer (BioRad). Commercially available antibodies used in this study are listed in Supplemental Table 2. Antibodies against LIN52 and phospho-S28-LIN52 were described in (Litovchick et al., 2011). Rabbit antibodies against DCAF7, FAM117B, LZTS1, LZTS2 and TROAP were custom samples provided by Bethyl.

### Glycerol Gradient Centrifugation

T98G cells were grown on two 150mm dishes and scraped using ice-cold PBS containing protease and phosphatase inhibitor cocktails (Calbiochem). The cells were collected by centrifugation and extracted using a buffer containing 10mM HEPES, 2mM MgCf_2_, 10mM KCl, 0.5% NP-40, 0.5mM EDTA, 150mM NaCl, H_2_O, 1mM DTT, protease and phosphatase inhibitors. For glycerol gradient analysis, 200 μL of whole cell lysate containing approximately 6 mg of protein was loaded on top of a pre-formed glycerol gradient in lysis buffer (5 ml, 5 - 45%). The sample was then centrifuged using SW55Ti rotor at 45,000 rpm at 4°C for 18 hours, after which 200 μL fractions were collected from the top of the gradient and analyzed by SDS PAGE electrophoresis using Criterion gels (BioRad) and Western Blotting.

### DSB repair Assay

DR-GFP reporter cell lines were established by transfecting the DR-GFP reporter construct (gift from Maria Jasin, Addgene plasmid # 26475 (Pierce et al., 1999)) into the control or DYRK1A-KO U-2 OS cells followed by puromycin selection as described in (Yakovlev, 2013). The cells stably expressing the DR-GFP reporter were infected with adenovirus to express I-SceI at MOI 50. To monitor the HRR efficiency, GFP positive cells were detected 48hr post-infection using flow cytometry as described in (Pierce et al., 1999; Yakovlev, 2013).

### Cell Cycle Analysis

U-2OS WT and DYRK1A-KO CRISPR cells were seeded in 10cm dishes at a density of 0.5X10^6^ - 1X10^6^.cells/dish. After 5Gy or 15Gy irradiation and incubation for 24hr, the cells were harvested and incubated with 0.05 mg/mL propidium iodide in 3.8 mM sodium citrate buffer containing 0.1% Triton X-100 and RNAse A solution (Sigma, R4642), and incubated at room temperature for 1 h. The cells (at least 10,000 per condition) were then analyzed using FACS Canto II flow cytometer (Becton Dickinson, San Jose, CA, USA),

### Statistical analysis and bioinformatic tools

For quantitation of cell-based experiments, 100 or more cells per conditions were typically scored. To calculate statistical significance, data from at least three biological replicates was analyzed using two-tailed Student’s t-test. For protein networks analysis, list of proteins detected in at least three out of four DYRK1A MudPIT analyses was analyzed using MetaScape web-based software (metascape.org) that integrates data from BioGrid (Chatr-Aryamontri et al., 2015) and other protein databases with custom datasets to build protein-protein interaction networks. To test the effect of DYRK1A expression on survival, level 3 gene expression data from TCGA summarized as RSEM values was obtained using TCGA2STAT R package v 1.2, along with corresponding clinical annotations. Data for breast cancer (BRCA) and lung adenocarcinoma (LUAD) cancers were obtained separately. To identify if the expression of DYRK1A affects survival in any specific clinical subgroup, subsets of patients annotated with specific clinical annotations were selected (e.g., “yes” or “no” in the “radiation” clinical annotation). The data was log2-transformed and analyzed using Kaplan-Meyer curves and Cox proportional hazard model. The expression of DYRK1A was analyzed for its effect on survival by separating patients into high/low expression subgroups. A modified approach from (Mihaly et al., 2013) was used to estimate the gene expression cutoff that best separates high/low expression subgroups with differential survival.

## Author Contributions

Conceptualization, L.L. and V.M; Methodology, L.L., V.M. and V.A.; Investigation, V.M., V.A., Sid.S., F.S., V.Y., Sel.S.,; Data Curation, Sel.S., L.A.F., L.L., M.D., V.A., V.M.; Writing -Original Draft, L.L. and V.M; Writing - Review & Editing, M.D., M.P.W., J.A.D.; Funding Acquisition, L.L.; Resources, M.P.W.; Supervision, L.L.

## Acknowledgements

Authors acknowledge Y. Skversky and S. Gruszecky for technical help, E. Ivanova and M. Tollenaere for help with cell imaging and G. D’Arcangelo for GFP-Dyrk1A constructs. We are grateful to N. Mailand for RNF169 constructs and for critical reading of the manuscript. This study was in part supported by N.I.H. R01CA188571 (L.L.). Proteomic studies were in part supported by the Stowers Institute for Biomedical Research. Services in support of this research project were generated by the VCU Massey Cancer Center Flow Cytometry Shared Resource as well as the VCU Microscopy Facility supported, in part, with funding from NIH-NCI Cancer Center Support Grant P30 CA016059.

